# Prefrontal Excitation/Inhibition Balance Supports Adolescent Enhancements in Circuit Signal to Noise Ratio

**DOI:** 10.1101/2024.08.15.608100

**Authors:** Shane D. McKeon, Maria I. Perica, Finnegan J. Calabro, Will Foran, Hoby Hetherington, Chan-Hong Moon, Beatriz Luna

**Author notes:** Corresponding author(s): Shane D. McKeon & Beatriz Luna. Lead Contact: Shane D. McKeon.

## Abstract

The development and refinement of neuronal circuitry allow for stabilized and efficient neural recruitment, supporting adult-like behavioral performance. During adolescence, the maturation of PFC is proposed to be a critical period (CP) for executive function, driven by a break in balance between glutamatergic excitation and GABAergic inhibition (E/I) neurotransmission. During CPs, cortical circuitry fine-tunes to improve information processing and reliable responses to stimuli, shifting from spontaneous to evoked activity, enhancing the SNR, and promoting neural synchronization. Harnessing 7T MR spectroscopy and EEG in a longitudinal cohort (N = 164, ages 10-32 years, 283 neuroimaging sessions), we outline associations between age-related changes in glutamate and GABA neurotransmitters and EEG measures of cortical SNR. We find developmental decreases in spontaneous activity and increases in cortical SNR during our auditory steady state task using 40 Hz stimuli. Decreases in spontaneous activity were associated with glutamate levels in DLPFC, while increases in cortical SNR were associated with more balanced Glu and GABA levels. These changes were associated with improvements in working memory performance. This study provides evidence of CP plasticity in the human PFC during adolescence, leading to stabilized circuitry that allows for the optimal recruitment and integration of multisensory input, resulting in improved executive function.

## 1. INTRODUCTION

Adolescence is a time of cognitive development that stabilizes in adulthood^2^, where the prefrontal cortex (PFC) is essential for cognitive processes and undergoes significant maturation, reflecting unique plasticity^1,3^. Postmortem studies in both animal and human models provide evidence of neurotransmitter changes involving glutamate and Gamma-Aminobutyric Acid (GABA)^4^, indicating a shift in E/I balance similar to critical period (CP) plasticity observed in sensory cortices earlier in development^1^. This shift includes alterations in parvalbumin (PV)-positive interneurons, a sub-type of inhibitory GABAergic interneuron known for its highly interconnected and fast-spiking activity, which enables them to synchronize and generate gamma oscillations observable via electroencephalography (EEG). Moreover, PV interneurons modulate their firing rates in response to excitatory input, serving as a crucial local gain control^11^ and contributing to the regulation of the E/I balance^1^. The fine-tuning of cortical circuitry during CPs enhances the circuit’s ability to produce consistent, reliable, and prompt responses to stimuli^1^. This transition involves a shift from predominantly spontaneous to evoked activity, which enhances the cortical SNR, facilitates neural population synchronization, and suppresses large asynchronous spontaneous activity^12–14^. This activity is thought to support executive functioning by stabilizing task-evoked activity, thus reducing performance variability on cognitive tasks for which accuracy and latency have been consistently shown to continue to improve into the twenties^2,5–10^. However, while evoked and spontaneous oscillations have been assessed, this shift in cortical SNR has not been studied *in vivo* in humans. Thus, this study aims to investigate cortical SNR using EEG and its associations with the E/I balance via MRSI measures of GABA and Glu, as a marker for CP plasticity during adolescence.

In early stages of neuronal development, spontaneous activity is prevalent and helps to shape neural circuit excitatory and inhibitory properties. For example, the influx of excitatory activity when light first hits the eye in the visual system helps to initiate a CP of circuit refinement^12,15^. The number of synapses in PFC are greater in adolescence than adulthood, which will be subsequently pruned^16,17^, resulting in increased excitatory over inhibitory activity. This hyperexcitable circuit function is thought to be important for the refinement of neuronal projections and neural wiring^18^. In response to the relative increase of excitatory function, the GABAergic inhibitory system undergoes significant maturation^19,20^ modulating neural activity by braking excitatory signaling. Thus, spontaneous activity during development may modulate excitatory and inhibitory synaptic efficacy^21^, thereby influencing the E/I balance throughout development and sculpting adult-like neuronal networks^22^. Animal models suggest that PFC GABAergic inhibitory circuitry undergoes significant alterations, including an increase in prefrontal PV+ interneurons during adolescence in the rat^23^ and non-human primate^24^. Furthermore, non-human primate and human postmortem studies have shown increases in PV interneurons^25^ and GABAA receptor α1 subunit expression^26^. Primarily expressed on PV interneurons, GABAA receptor α1 subunits support fast synaptic inhibition^27^ and synaptic plasticity^28^. In conjunction, excitatory pyramidal PFC neurons undergo significant pruning across the adolescent period^29^. These findings provide evidence suggesting CP plasticity in PFC akin to CPs observed in sensory systems, such as the development of PV+ interneuron circuitry and excitatory circuit maturation^30,31^, which facilitate experience-driven CP plasticity^28,32^.

The interplay between PV+ interneurons, in particular basket cells, that innervate on the soma of pyramidal neurons, generate gamma oscillations^33–35^. The regulation of which occurs at both the single neuron level, regulated by the number of excitatory and inhibitory synapses on cortical pyramidal neurons, and the large-scale circuitry level, where numerous homeostatic and developmental processes help maintain the E/I balance across long timescales^36^. These two levels, while independent, work in conjunction as stimulated neurons in one population propagate other neurons across multiple cortical layers, until sufficient inhibition is reached to maintain balance^37,38^. Coupled together, both levels contribute to the appropriate modulation of the inhibition excitation activity needed to generate gamma oscillations^39^, allow for efficient information transmission and gating^40^, network computation^41^, and higher order cognitive functions, such as working memory^42^.

Developmental changes in spontaneous and evoked activity have been studied via the auditory steady state response (ASSR), which uses a train of auditory clicks delivered at high frequencies, typically at 20, 30 or 40Hz, that have been demonstrated to elicit robust oscillations that can be noninvasively measured through EEG. This phenomenon, known as entrainment, is driven by a phase adjustment of the underlying oscillations to match the stimulus frequency and an increase in amplitude at the neural population level as more neurons become aligned to the stimuli^43,44^. This is thought to be propelled by a circuit’s preferred resonance frequency at which the auditory stimuli induces the most entrainment^45,46^, and is referred to as the evoked activity. In contrast, spontaneous power of gamma oscillations is non-stimulus locked, allowing the parsing of task-locked from non-locked activity to infer relative levels of spontaneous and evoked activity and, consequently, cortical SNR. Previous studies have demonstrated that the greatest and most reliable evoked response is yielded by a 40 Hz stimuli^45,47^, and has been historically used to assess the ability to generate gamma-band (30-80 Hz) activity^34,48–62^. Furthermore, animal models have shown the 40 Hz ASSR is maintained by N-methyl-D-aspartate (NMDA) receptors on pyramidal neurons and GABA systems^63–65^, demonstrated via pharmacological studies using NMDA antagonist ketamine^66,67^ and the GABAA agonist muscimol^63^, where power in the 40 Hz ASSR increases.

While the ASSR has been observed in auditory-responsive regions, including those in the temporal and parietal lobes^68–71^, it has also been observed in PFC regions when presented with a 40 Hz stimulation^68,72–78^. Previous studies interrogating the underlying source of the ASSR have looked at the tuning characteristics of individual electrodes and found different characteristics between parietal and frontal cortex, suggesting the auditory response is occurring in parallel in different, pathways^75^. One potential explanation is that auditory entrainment requires top-down attentional modulation which may be maintained by the frontoparietal attention network^79,80^.

Given that the ASSR is generated by cycling between GABAergic inhibition and rebound excitation, is observed in the frontal cortex, and that the GABAergic system is undergoing significant maturation during adolescence, we hypothesize that the ASSR may reflect underlying consequences of E/I balance in frontal regions. Previous developmental studies have found increases in evoked power from 40 Hz auditory stimuli from ages 5-52^81^ and ages 19-45^82^, revealing an inverted-U trajectory. This pattern is thought to reflect the interplay between opposing developmental effects: the increases in α1 GABA receptors promoting gamma activity, increasing the evoked activity, and the later synaptic pruning of pyramidal cell excitation^34^, consequently decreasing the evoked activity. Furthermore, previous studies have found resting state power, or spontaneous power, to decrease across all frequency bands from early to late childhood^83^, supporting evoked activity, thus increasing cortical SNR, that may be reflecting circuitry plasticity^1,13,84^.

Given that inhibitory circuitry maturation regains E/I balance, and increases cortical SNR during animal model CPs^14^, we hypothesized cortical SNR would increase with age, driven by increases in evoked power and decreases in spontaneous activity, as measured by the ASSR task. In this study, we collected a large, multimodal, longitudinal dataset with EEG and 7T Magnetic Resonance Spectroscopic Imaging (MRSI) of PFC, to investigate developmental changes in cortical SNR and its association with the E/I balance. We have previously showed neuroimaging evidence for increases in Glu/GABA balance into adulthood in PFC^85,86^. Thus, we hypothesized that age related increases in E/I balance would be associated with increases in cortical SNR, derived from a 40 Hz stimuli auditory steady state task. In line with this hypothesis, we found developmental increases in PFC cortical SNR, driven by decreases in spontaneous activity, where SNR was found to be significantly associated with increases in PFC E/I balance. Finally, we show that increases in cortical SNR and decreases in spontaneous activity were associated with age related behavioral improvements in accuracy and latency variability in the memory guided saccade task. Taken together, this study provides *in vivo* evidence for CP plasticity mechanisms, i.e. increases in the E/I balance driving increases in cortical SNR, in the PFC through adolescence supporting stable and efficient circuitry and ultimately cognitive development.

## 2. METHODS

### 2.1 Participants

Data was collected on 164 participants (87 assigned female at birth), between 10-32 years of age. Participants were recruited as part of an accelerated cohort design with up to 3 visits at approximately 18mo intervals. Each time point consisted of three visits: a behavioral (in-lab) session, a 7T MRI scan, and an EEG session, typically occurring on different days within 1-2 weeks, for a total of 347 visits. Following data quality control and exclusion criteria, as described below, the final dataset included 283 sessions. Participants were recruited from the greater Pittsburgh area and were excluded if they had a history of loss of consciousness due to a head injury, non-correctable vision problems, learning disabilities, a history of substance abuse, or a history of major psychiatric or neurologic conditions in themselves or a first-degree relative. Patients were also excluded if any MRI contradictions were reported, including but not limited to, non-removable metal in their body. Participants or the parents of minors gave informed consent with those under 18 years of age providing assent. Participants received payment for their participation. All experimental procedures were approved by the University of Pittsburgh Institutional Review Board and complied with the Code of Ethics of the World Medical Association (Declaration of Helsinki, 1964).

### 2.2 Data Acquisition and Preprocessing

#### 2.2.1 Electrophysiological (EEG) Data

Concurrent EOG (electrooculogram) and high-impedance EEG was recorded using a Biosemi ActiveTwo 64-channel EEG system located in the PWPIC Psychophysiology Laboratory and the University of Pittsburgh. EEG sessions were conducted in an electromagnetically shielded room while stimuli were presented by a computer approximately 80 cm away from participants. EEG data was collected during our auditory steady state task where participants are presented with a 20, 30, and 40 Hz click train. Initial auditory data was sampled at 1024 Hz and resampled at 512 Hz during preprocessing. Scalp electrodes were referenced to external electrodes corresponding to the mastoids due to its proximity to the scalp and low signal recording. An initial bandpass filter was set to 0.5 – 70 Hz. Data were preprocessed using a revised processing pipeline compatible with EEGLAB^87^, which removed flatline channels (maximum tolerated flatline duration: 8 seconds), low-frequency drifts, noisy channels (defined as more than 5 standard deviations from the average channel signal), short spontaneous bursts, and incomplete segments of data. Deleted channels were replaced with interpolated data from surrounding electrodes. The resulting data was referenced to the average reference. As a final preprocessing step, independent component analysis (ICA) was performed to identify eye-blink artifacts and remove their contribution to the data.

#### 2.2.2 Magnetic Resonance Spectroscopy

MRSI methods have been previously reported in Perica et al. (2022)^86^. Briefly, data were acquired at the University of Pittsburgh Medical Center Magnetic Resonance Research Center using a Siemens 7T scanner. Structural images were acquired using an MP2RAGE sequence (1 mm isotropic resolution, TR/TE/flip angle 1/ flip angle 2: 6000 ms/2.47 ms/40/50). MRSI including GABA and glutamate were acquired using a J-refocused spectroscopic imaging sequence (TE/TR = 2 ×17/1500 ms) and water suppression was performed using a broad band semi-selective refocusing pulse and frequency selective inversion recovery^88^. Radiofrequency (RF) based outer volume suppression was used to minimize interference in the signal from extracerebral tissue^89^. An 8 × 2 1H transceiver array using 8 independent RF channels was used to acquire data. High order shimming was used to optimize homogeneity of the B0 magnetic field. The 2D CSI oblique axial slice was acquired with conventional rectangular phase encoding of 24 × 24 over a FOV of 216 × 216 mm (10 mm thick, 0.9 ×0.9 ×1.0 cm nominal resolution), and was positioned to include Brodmann Area 9 and pass through the thalamus. Methodologcial representation can be seen in Figure *3*A.

#### 2.2.3 Auditory Steady State Task

Participants were presented with a blank screen and an auditory train of clicks in stereo via over-ear headphones. The stimulus was presented in blocks of 150 trials, each of which consisted of a 500ms click train followed by 605ms of silence. Blocks contained either ten 20Hz clicks, fifteen 30 Hz clicks, or twenty 40 Hz clicks. Block order was randomly assigned (e.g., 30-40-20) for each participant at each session. The task was written in and executed by NBS Presentation software.

#### 2.2.4 Memory Guided Saccade Task

As reported in our previous studies^10^, participants performed a memory guided saccade (MGS) task to assess working memory performance during the EEG session. Participants performed 3 runs of the MGS task, each containing 20 trials comprised of a visual guided saccade (VGS) to a peripheral cue, variable delay epoch (6-10 sec) where the participant must maintain the location of the peripheral target in working memory, followed by a memory guided saccade (MGS) to the remembered location. Task performance was assessed based on horizontal electrooculogram (hEOG) channels recorded from facial muscles (see acquisition details below). These were used to calculate VGS & MGS response latencies, as the time difference between the beginning of the recall epoch and the initiation of the VGS and MGS eye movements respectively, and saccadic accuracy, measured as the closest stable fixation point during the recall period to the fixated location during the initial visually guided fixation epoch. Visual representation of the task can be seen in Figure 4A.

### 2.3 Data Analysis

#### 2.3.1 EEG Analyses

##### 2.3.1.1 Cortical Signal-to-Noise Ratio (SNR)

Cortical signal-to-noise ratio (SNR) was computed by calculating the evoked (stimulus-locked) and total power during the 20, 30, and 40Hz click train frequency conditions. Prestimulus values (-200 to 0ms) were extracted to provide a baseline reference. A task epoch was then defined over the first 800ms following stimulus onset, and the auditory steady state response (ASSR) was computed for each electrode based on methods based on previous literature^50,61,90,91^. Briefly, a Fast Fourier Transform (FFT) was applied to each single-trial epoch, resulting in per-trial power spectra. Power spectra were then averaged across trials to compute total power as a function of frequency. Evoked power was then derived by averaging the single trial time courses and calculating the resulting power spectra using FFT. By averaging across trials, only activity which was consistently timed relative to the stimulus onset remained, providing an estimate of the stimulus-evoked power. Spontaneous power was then derived by subtracting the evoked power from the total power. Finally, SNR was calculated as the ratio of evoked power to spontaneous power. A methodological representation of the method can be seen in *Figure 1*.

**Figure 1.**
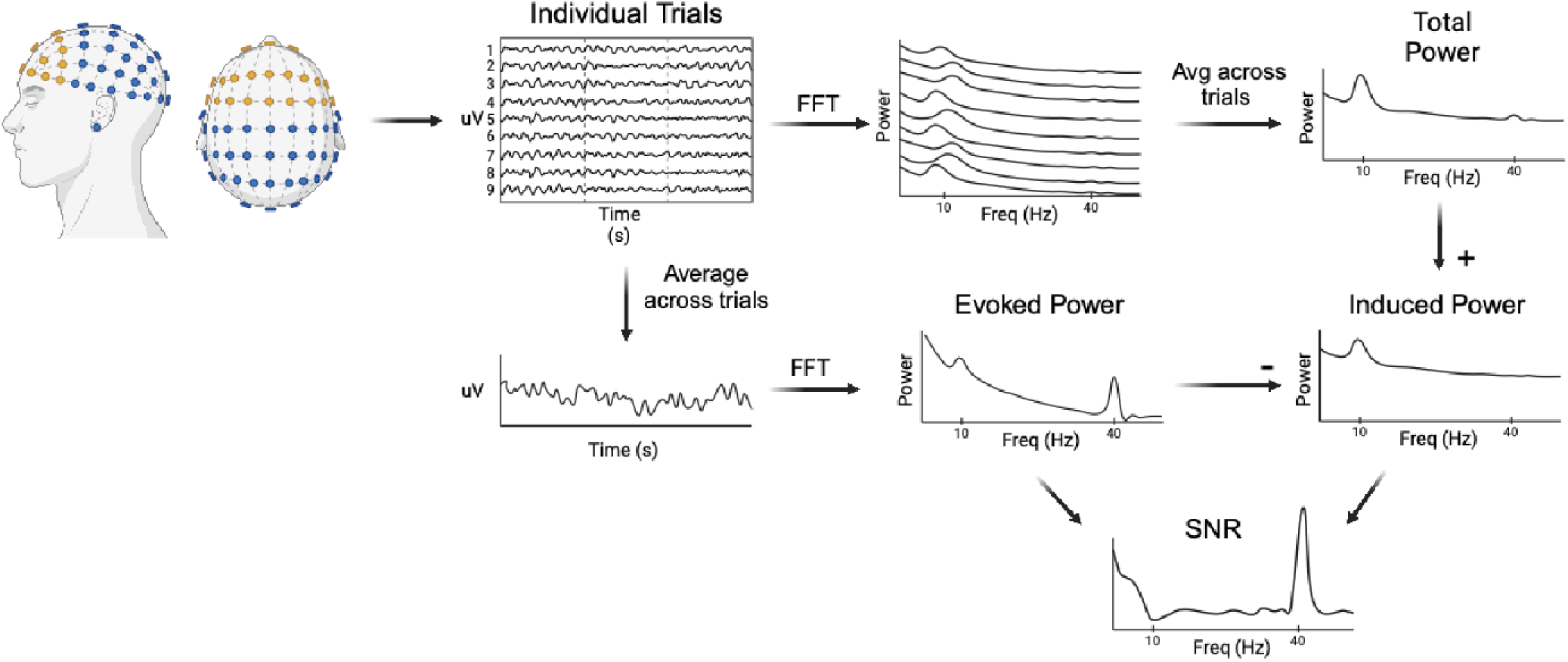
Methodological representation for evoked power, spontaneous power, and cortical SNR.

##### 2.3.1.2 Dimensionality Reduction using PCA

Prior work has shown that ASSR is maximal at the frontal central electrodes Fz and FCz and then disperses amongst the frontal central electrodes surrounding it^59,61,73,92,93^. To account for this spatial signal spread, we performed a principle component analysis (PCA) to reduce the dimensionality of the data for each of the total, evoked, and spontaneous power, as well as SNR. Due to both prior work and our hypotheses regard prefrontal cortex plasticity, the PCA was restricted to frontal electrodes, that is, F3, F5, F7, F1, F2, F4, F6, F8, AFz, AF1, AF2, Fp1, Fp2, Fz, AF5, AF6 (per the 10-20 international system naming convention). In each case, the first principal component (PC) captured the bulk for the signal variance and was used for subsequent analyses (Figure *2*A).

**Figure 2.**
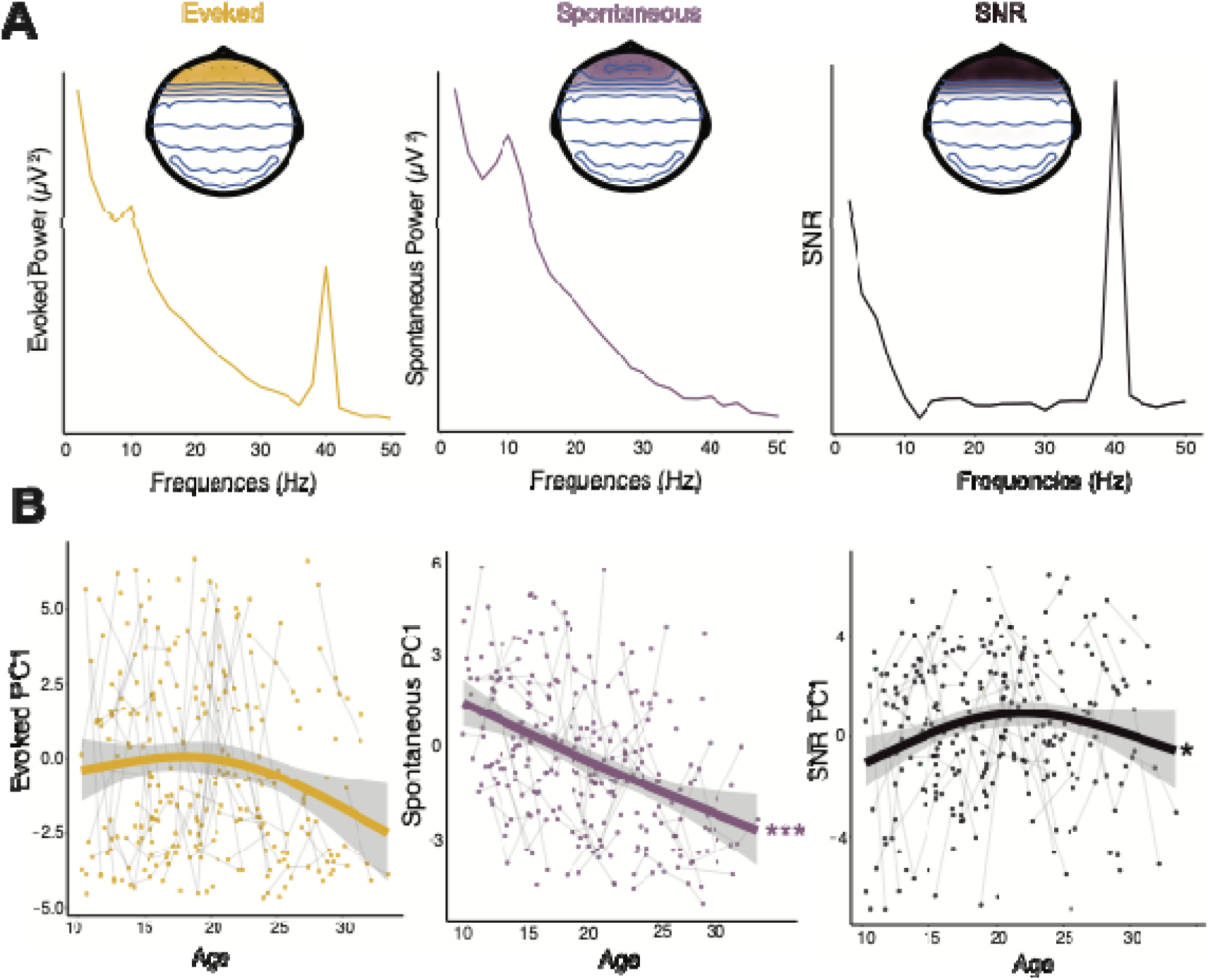
**A**. Brain plots represent frontal ROI used for PCA. (Left) Evoked power by frequency. (Middle) Spontaneous power vs frequency. (Right) Cortical SNR vs frequency. **B.** (Left) Evoked activity PC1 values vs age (NS = p = 0.137). (Middle) Spontaneous activity PC1 values vs age. (Right) Cortical SNR PC1 values vs age. (* p < 0.01; ** p < 0.001; *** p < 0.0001).

Analysis details have been previously reported in Perica et al. (2022)^86^. Briefly, a 2D CSI oblique axial slice was acquired (TR/TE/flip angle: 2180 ms, 23 ms, 7°, voxel size = 2.0x2.0x2.0 mm), as outlined in Perica et al. (2022), to ensure that the dorsolateral prefrontal cortex (DLPFC) was included and angled to pass through the thalamus. Regions of interest included right and left DLPFC due to its role in working memory. LCModel was used to obtain estimates of individual metabolites^94^, including creatine (Cre), γ-aminobutyric acid (GABA), and glutamate (Glu), among others. Glu-GABA asymmetry was calculated by taking the absolute value residual of the linear model of the association between Glu/Cr and GABA/Cr. This measure captures the correlation of Glu and GABA levels thus controlling for the confound of age related variability in Glu/GABA ratio^95,96^. Based on a Kolmogorov-Smirnov test of normality, the Glu-GABA asymmetry was found to be non-normal (p = 4.39e-07). Thus, a square root correction was applied to improve normality (resulting p = 0.052).

#### 2.3.2 Behavioral Analyses

To characterize development of behavioral measures from the MGS task, including accuracy and response latency, we fit age-related performance effects using generalized additive mixed models (GAMMs) using the R package mgcv^97^. Preliminary outlier detection was conducted on a trial level basis. Express saccades, in which eye movements were initiated within the first 100ms after task onset were excluded, since these are believed to be primarily driven by subcortical systems without involving cortical processing of the visual cue^98^. Position error measures greater than 23 degrees from the target were excluded as implausible since they exceeded the width of the screen. The remaining trials for each participant were combined, creating average performance measures and trial-by-trial variability measures for each participant. A separate model was performed for each of the behavioral measurements: MGS accuracy, MGS latency, as well as the trial-by-trial variability of each measure. Behavioral data from this task has been previously presented in McKeon et al. (2023)^99^.

#### 2.3.3 Statistical Analyses

A threshold of 2 standard deviations from the mean was used to exclude statistical outliers for each electrode before the PCA. Due to missing data from outlier detection on each electrode, we imputed the missing data, using RStudio package mice^100^, for each subject so not to lose subjects who did not have complete data for all channels. Individual outliers were then detected using 2 standard deviations from the mean for each PC value for the evoked, spontaneous and SNR measures, as well as the MRSI derived measures, Glu, GABA, and Glu/GABA asymmetry.

To assess developmental trajectories of cortical SNR activity, we implemented GAMMs on the first principal component, PC1, of evoked power, spontaneous power, and SNR, including random intercepts estimated for each participant. Regression splines were implemented (4 degrees of freedom) to assess linear and non-linear effects^97,101^. Auditory measures that were found to significantly change across adolescence were then used to test for associations with our MRSI measures, glutamate (Glu), GABA, and Glu GABA Asymmetry using linear mixed effect models (lmer function, lme4 package in Rstudio^102^). We first tested for significant main effects of the auditory measure on the MRSI parameter while controlling for age and hemisphere (left or right DLPFC). We additionally tested for auditory measure-by-age interactions while controlling for hemisphere. We then investigated whether our auditory measures had significant associations with our working memory measures (accuracy, accuracy trial variability, response latency, response latency variability) using linear mixed effect models (lmer function, lme4 package in Rstudio^102^).

## 3. RESULTS

### 3.4.1 Cortical SNR increases across adolescence

Auditory activity was first extracted to assess the validity of our task. Evoked activity (Figure *2*A Left) was found to peak at the frequencies corresponding to the auditory stimuli (i.e. 20, 30 and 40Hz), and in the case of the 20 Hz condition, a peak was also seen in the harmonic 40 Hz band (first harmonic). Spontaneous activity (Figure *2*A Middle), derived from the difference between total and evoked activity, did not show these peaks at the corresponding frequencies, supporting the applied method for separating task-evoked from spontaneous activity. Cortical SNR, derived as the ratio of evoked activity/spontaneous activity, showed peaks at the corresponding frequencies to auditory stimuli and their related harmonic frequencies (Figure *2*A Right). To assess the developmental changes for each measure across adolescence, we tested for associations between age and each measure’s first principle component. Evoked activity did not change with age (F = 1.45, p = 0.137; (Figure *2*B Left), while spontaneous activity showed significant decreases with age (F = 37.53, p < 2e-16; Figure *2*B Middle). Consequently, cortical SNR was found to increase through adolescence (F = 4.59, p = 0.045; Figure *2*B Right).

#### 3.4.2 Associations between cortical SNR and MRSI

To test if cortical SNR ratio is associated with MRSI-derived measures of E/I balance, we assessed relationships between EEG-derived auditory measures and Glu, GABA, and Glu-GABA asymmetry. All statistics were conducted controlling for age and hemisphere (left vs right DLPFC). Greater Glu was associated with less spontaneous activity (β = -0.015, t = -2.28, p = 0.023; Figure *3*B) while age-by-glutamate interaction was not significant (β = -0.001, t = -1.41, p = 0.16). Glutamate did not have a significant relation with cortical SNR (β = 0.006, t = 1.24, p = 0.66). Greater Glu-GABA asymmetry was associated with less PFC SNR (β = - 0.007, t = -2.38, p = 0.017; Figure *3*C) but did not have a significant age-by-Glu-GABA asymmetry interaction (β = 0.12, t = 1.02, p = 0.31). Further analysis showed that Glu-GABA asymmetry was not significantly associated with spontaneous activity (β = 0.005, t = 1.175, p = 0.72).

**Figure 3.**
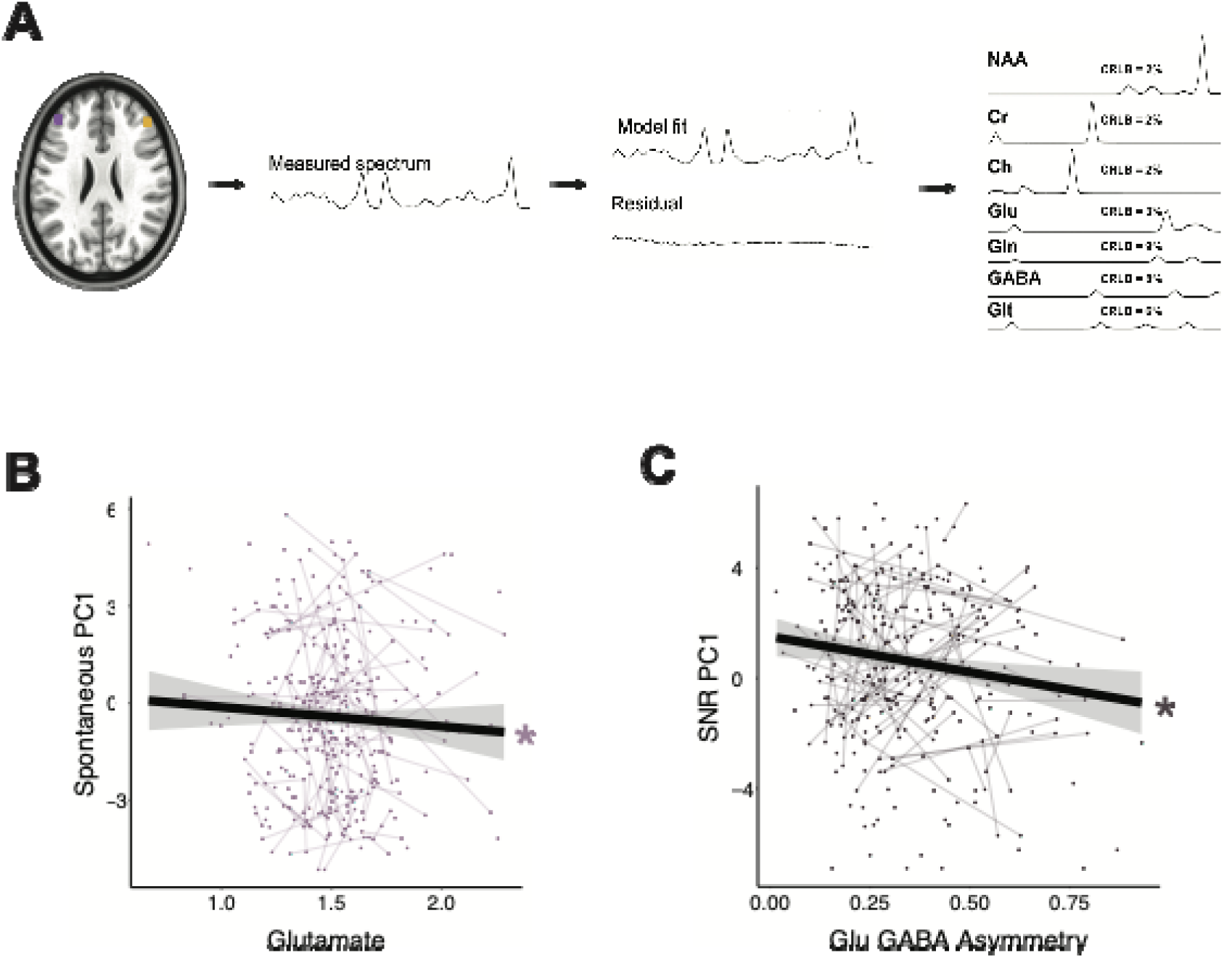
**A.** Methodological representation of MRSI to measure glutamate and GABA. **B.** Spontaneous activity PC1 values vs glutamate. **C.** Cortical SNR PC1 values vs Glu GABA Asymmetry. (* p < 0.01).

### 3.4.3 Associations between cortical SNR and working memory

An essential aspect of adolescent development involves the ongoing enhancement of executive functions, such as working memory, believed to rely on balance within excitatory/inhibitory (E/I) circuitry^42^ and signal-to-noise ratio (SNR) of cortical circuits^31,103^. To assess whether SNR was related to improvements in executive function, we investigated the associations between cortical SNR and working memory, based on performance on the memory guided saccade (MGS) task. We found that greater PFC SNR was associated with better performance accuracy (measured as degrees away from the intended target) (β = -0.52, t =-2.56, p = 0.011; Figure 4C Right). Neither evoked (β = -0.25, t = -1.09, p = 0.27) or spontaneous (β = 0.21, t = 1.41, p = 0.16) activity was significantly associated with performance accuracy. Meanwhile, we found the trial-by-trial response latency variability increased with spontaneous activity (β = 14.7, t = 2.85, p = 0.005; Figure 4D Middle) and with decreases in PFC SNR (β = -16.34, t = -2.23, p = 0.03; Figure 4D Right). Spontaneous activity did not have a significant age-by-latency variability interaction (β = 0.93, t = 1.07, p = 0.28) nor did PFC SNR (β = -0.47, t = -0.38, p = 0.68). (Additional behavioral measures, accuracy trial variability and MGS response latency can be found in Supplement Figure 1. Additional statistics can be found in Supplement Table 1).

**Figure 4.**
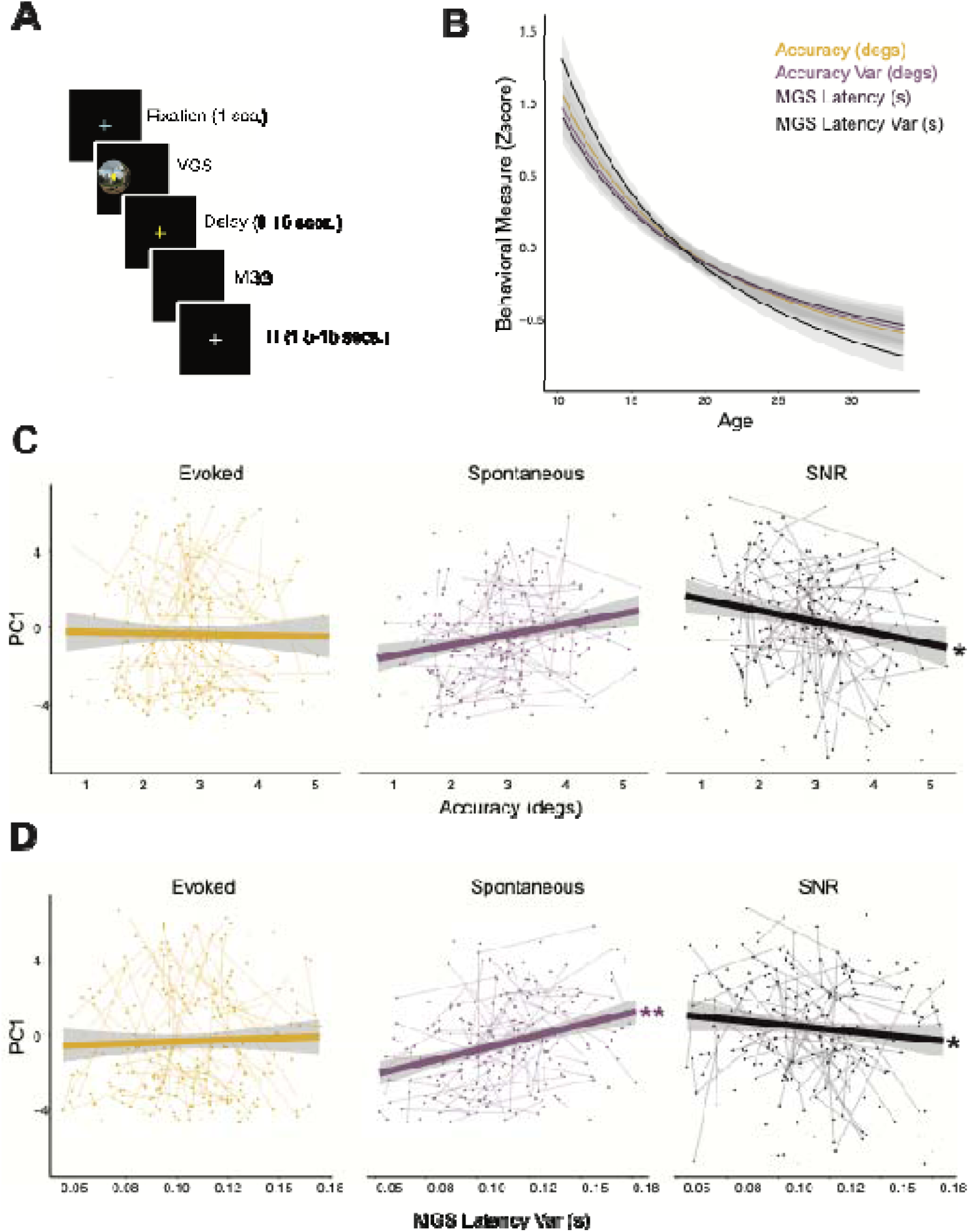
**A.** Memory guided saccade task used to assess working memory. **B.** Working memory performance measures (accuracy, accuracy variability, response latency, and response latency variability) across age. Previously reported in McKeon, 2024^85^. **C**. Auditory measures, evoked activity (left) (NS p=0.27), spontaneous activity (middle) (NS p=0.16), and cortical SNR (right) vs accuracy (measured as degrees away from intended target). **D.** Auditory measures, evoked activity (left), spontaneous activity (middle), and cortical SNR (right) vs trial-by-trial variable of response latency (measured as degrees away from intended target). (** p < 0.001; * p < 0.01).

## 4. DISCUSSION

This study aimed to investigate PFC SNR using EEG and its associations with the E/I balance via MRSI measures of GABA and Glu, as a mechanism of CP plasticity during adolescence. Previous animal studies investigating sensory cortex CPs has shown that maturation of the GABAergic inhibitory system supports the suppression of spontaneous, asynchronous activity, in favor of evoked, synchronous activity, increasing the cortical SNR^12–14^. Consistent with these CP mechanisms, we found that PFC cortical SNR, during 40 Hz auditory steady state task, increases across adolescence, driven by significant decreases in spontaneous activity. Furthermore, we found increases in PFC SNR to be associated with increases in the DLPFC Glu/GABA balance. Post-hoc analyses suggested this association is driven by the significant relationship between increases in spontaneous activity and decreases in glutamate. To interrogate the developmental effects of PFC SNR on behavior, we assessed working memory via a memory guided saccade (MGS) task, results of which have previously been reported^10,85^. Here, we found increases in PFC SNR to be correlated with increases in performance accuracy suggesting that fidelity of the mnemonic information is supported by optimal SNR. While decreases in spontaneous activity, and increases in PFC SNR, were found to be associated with decreases in performance latency variability, suggesting that performance stability is supported by both high SNR and associated decreases in spontaneous neural function. Taken together, these results suggest developmental increases in PFC SNR, driven by decreases in spontaneous activity, may be due to developmental increases in Glu/GABA balance, and support the development of higher order cognitive functions through adolescence into adulthood.

Previous work has shown that the auditory steady state task elicits robust gamma oscillations driven by 40Hz stimulus clicks, thought to be driven by GABAA receptors^104^ via cycling through inhibition and rebound excitation^63,105^. Furthermore, the ASSR is strongest at central electrodes Fz and FCz, with activity dispersing throughout the frontal electrodes^59,61,73,92,93^. Given that the GABAergic system undergoes significant maturation through adolescence in the PFC^23–25,104,106,107^, we hypothesized that the frontal ASSR should change developmentally. Consistent with this hypothesis, previous developmental studies have found increases in evoked power from 40 Hz auditory stimuli from ages 5-52yo^81^ and 19-45yo^82^, with a study spanning ages 8-22 identifying an inverted-U trajectory^108^. These studies however found the most significant changes in young childhood, roughly ages 5-10 years old, in contrast to our older cohort of 10-32 years of age, suggesting that evoked activity may mature in childhood. These patterns of evoked power across development are thought to reflect the interplay between opposing developmental effects: the increases in α1 GABA receptors promoting gamma activity, thus increasing evoked activity, and the later synaptic pruning of pyramidal cell excitation^34^, which in PFC, begins in childhood continuing through adolescence^109^, thus decreasing the evoked activity.

Spontaneous activity is conceptually similar to brain activity during resting state, that is, activity that is not locked to any task stimuli. Trial-by-trial power variations during tasks^110,111^, and non-stimulus-locked activation during auditory steady state^61^ decrease across adolescence in all frequency bands^10,85,112–115^. Here, we sought to assess spontaneous activity, sometimes referred to as induced activity, that is non-stimulus-locked, across adolescence, during a 40 Hz auditory steady state task. Excitatory activity is thought to be driven by NMDA-mediated activation on PV positive interneurons, triggering postsynaptic potentials, and generating gamma oscillations^116–118^. Here, we found highly significant decreases in spontaneous activity across age, which may be indicative of PV+ interneuron maturation that has been found to suppress spontaneous firing^12–14^. Furthermore, NMDA subunit NR2B density, which increases in transmodal cortices^119,120^ at the beginning of a critical period, begins to decrease as the critical period closes^121^, which may further explain the decrease in spontaneous firing, constrained by the shortening of the time course of excitatory synapses. This switch from spontaneous to evoked power has been suggested to increase cortical SNR across adolescence, due to PFC PV positive basket cells ability to innervate pyramidal neurons on their soma allow them to synchronize activity between cortical networks^30,31^. We then calculated cortical SNR as the ratio between evoked activity and spontaneous activity and found significant increases across adolescence. These results suggest PFC is undergoing similar mechanisms as sensory system CPs, but in adolescence such as the maturation of PV+ interneuron circuitry maturation^30,31^, and excitatory circuitry maturation, promoting CP plasticity^28,32^.

In our previous work we have shown developmental decreases in MRSI derived levels of glutamate and increases in the Glu/GABA balance^85^ throughout adolescence into adulthood. Here, we sought to investigate whether these measures of the E/I balance would be associated with our EEG derived measures of cortical SNR. Indeed, we find increases in Glu/GABA balance, measured as a decrease in the Glu/GABA asymmetry, to be significantly associated with increases in cortical PFC SNR. Post-hoc analyses showed that this relationship may be underlined by the significant association, when controlling for age, where ultimately decreases in spontaneous activity in PFC are associated with increases in glutamate. While this result may seem counterintuitive, it may be representing the contrasting developmental mechanisms of strengthening synaptic connections, network integration, and sustained excitatory activity^76,122,123^, all while the cortex undergoes significant synaptic pruning^109,124^. Furthermore, the NMDA subunit change during adolescence^119,120^ first supports excitatory activity^76,122,123^ via glutamate, but as the CP begins to close, the subunit density decreases and maintains a low level in adulthood, thus contributing to decreases in excitation^121^. These complex and contrasting developmental mechanisms need to be further investigated.

A key part of adolescent development is the continued refinement of executive functions^2^, including working memory, thought to be supported by increasing E/I balance^42^, which in turn supports a high cortical SNR^31,103^. Accuracy and latency in working memory are unique components underlying executive function, where accuracy reflects the fidelity of the mnemonic information being kept in working memory, while latency reflects the speed of cognitive information processing^125,126^. Further, the DLPFC, the region of interest used in our MRSI data, is a region known to be involved in visual spatial memory^127^. As we previously reported^85^, working memory accuracy, trial-by-trial accuracy variability, response latency, and trial-by-trial response latency show improvements into adulthood. Here, we compared these behavioral results to our measures of evoked, spontaneous activity, and PFC SNR. We found performance accuracy improved with increasing cortical SNR. We also found decreases in trial-by-trial response latency variability to be associated with decreases in spontaneous activity and increases in cortical SNR. These results suggest that age-related improvement in accuracy may be due to the E/I balance maximizing signal efficiency of neural responses via local gain control on excitatory circuitry^33,128^ while increases in the speed of generating a response may reflect stabilization of optimal processing.

CP mechanisms result in neural circuitry that has high SNR to a given input and responds with a synchronous, consistent output sending activity with high SNR to downstream circuits^1^. To perform higher order cognitive functions, such as in working memory, areas that integrate sensory information, such as the PFC, require stable inputs. During adolescence, sensory systems are providing high SNR output to downstream circuits that support task performance, like the PFC. Thus, our results suggest that with maturation the PFC integrates multisensory input into stable and consistent activity, reflected in increasing cortical SNR and greater E/I balance, resulting in enhanced ability to maintain accurate information online. This may be achieved through maximizing signal efficiency of neural responses via local gain control on excitatory circuitry^33,128^. Furthermore, high SNR drives consistent stimulus-evoked activity, and suppresses spontaneous activity. Thus, response latency variability improvements being associated with reductions in spontaneous activity may suggest that activity stabilizes and becomes more reliable as synaptic pruning establishes optimal circuitry.

Together, these results provide *in vivo* evidence that PFC SNR increases as the E/I circuitry becomes balanced, supporting the refinement and integration of multipart sensory input into a stable and reliable circuitry that can perform complex executive functioning. Importantly, CP mechanisms can inform the basis of refinements in higher-order cognition and provide context for structural and functional maturation during adolescence^1^. Furthermore, these results are in line with a model of adolescent PFC CP plasticity^129^, as well as, build off our previous work on the aperiodic component as a measure of E/I balance and CP plasticity during adolescence^85^. Importantly, these results provide possible mechanisms for plasticity in normative development but also in impaired development, such as in psychopathology, which predominantly emerges in adolescence (e.g., psychosis, mood disorders, addiction)^130,131^, is associated with limitations in cognition^54,82,132^, and with impairments in excitatory and inhibitory processes^54,104,133–135^.

## Acknowledgements

We thank the University of Pittsburgh Clinical and Translational Science Institute (CTSI) for support in recruiting participants, as well as their support by the National Institutes of Health through grant number <u>UL1TR001857</u>. We thank Matthew Missar, Laurie Thompson, Alyssa Famalette, Vivian Lallo, Piya Verma, and Angela Martinez for their work involving our data collection.

## References

1. Larsen, B. & Luna, B. Adolescence as a neurobiological critical period for the development of higher-order cognition. Neurosci Biobehav Rev 94, 179–195 (2018).

2. Tervo-Clemmens, B. et al. A canonical trajectory of executive function maturation from adolescence to adulthood. Nat Commun 14, 6922 (2023).

3. Fuster, J. M. Frontal lobe and cognitive development. J Neurocytol 31, 373–385 (2002).

4. Kilb, W. Development of the GABAergic System from Birth to Adolescence. Neuroscientist 18, 613–630 (2012).

5. Montez, D. F., Calabro, F. J. & Luna, B. The expression of established cognitive brain states stabilizes with working memory development. Elife 6, (2017).

6. Simmonds, D. J., Hallquist, M. N. & Luna, B. Protracted development of executive and mnemonic brain systems underlying working memory in adolescence: A longitudinal fMRI study. Neuroimage 157, 695–704 (2017).

7. Alloway, T. P., Gathercole, S. E. & Pickering, S. J. Verbal and Visuospatial Short-Term and Working Memory in Children: Are They Separable? Child Development 77, 1698–1716 (2006).

8. Luna, B., Garver, K. E., Urban, T. A., Lazar, N. A. & Sweeney, J. A. Maturation of cognitive processes from late childhood to adulthood. Child Dev 75, 1357–1372 (2004).

9. Geier, C. F., Garver, K., Terwilliger, R. & Luna, B. Development of Working Memory Maintenance. J Neurophysiol 101, 84–99 (2009).

10. McKeon, S. D. et al. Age-related differences in transient gamma band activity during working memory maintenance through adolescence. NeuroImage 120112 (2023) doi:10.1016/j.neuroimage.2023.120112.

11. Scholl, B., Pattadkal, J. J., Dilly, G. A., Priebe, N. J. & Zemelman, B. V. Local Integration Accounts for Weak Selectivity of Mouse Neocortical Parvalbumin Interneurons. Neuron 87, 424–436 (2015).

12. Hensch, T. K. Critical period plasticity in local cortical circuits. Nat Rev Neurosci 6, 877–888 (2005).

13. Toyoizumi, T. et al. A theory of the transition to critical period plasticity: inhibition selectively suppresses spontaneous activity. Neuron 80, 51–63 (2013).

14. Hensch, T. K. & Fagiolini, M. Excitatory–inhibitory balance and critical period plasticity in developing visual cortex. Progress in Brain Research 147, 115–124 (2005).

15. Wiesel, T. N. & Hubel, D. H. SINGLE-CELL RESPONSES IN STRIATE CORTEX OF KITTENS DEPRIVED OF VISION IN ONE EYE. J. Neurophysiol. 26, 1003–1017 (1963).

16. Kolb, B. & Gibb, R. Brain Plasticity and Behaviour in the Developing Brain. J Can Acad Child Adolesc Psychiatry 20, 265–276 (2011).

17. Huttenlocher, P. R. Synaptic density in human frontal cortex - developmental changes and effects of aging. Brain Res 163, 195–205 (1979).

18. Mazzoni, A. et al. On the Dynamics of the Spontaneous Activity in Neuronal Networks. PLOS ONE 2, e439 (2007).

19. Wu, C. & Sun, D. GABA receptors in brain development, function, and injury. Metab Brain Dis 30, 367–379 (2015).

20. Li, J., Chen, L., Guo, F. & Han, X. The Effects of GABAergic System under Cerebral Ischemia: Spotlight on Cognitive Function. Neural Plasticity 2020, e8856722 (2020).

21. Gonzalez-Islas, C. & Wenner, P. Spontaneous network activity in the embryonic spinal cord regulates AMPAergic and GABAergic synaptic strength. Neuron 49, 563–575 (2006).

22. Haider, B., Duque, A., Hasenstaub, A. R. & McCormick, D. A. Neocortical Network Activity In Vivo Is Generated through a Dynamic Balance of Excitation and Inhibition. J. Neurosci. 26, 4535–4545 (2006).

23. Caballero, A., Flores-Barrera, E., Cass, D. K. & Tseng, K. Y. Differential regulation of parvalbumin and calretinin interneurons in the prefrontal cortex during adolescence. Brain Struct Funct 219, 395–406 (2014).

24. Hoftman, G. D. et al. Altered cortical expression of GABA-related genes in schizophrenia: illness progression vs developmental disturbance. Schizophr Bull 41, 180–191 (2015).

25. Fung, S. J. et al. Expression of interneuron markers in the dorsolateral prefrontal cortex of the developing human and in schizophrenia. American Journal of Psychiatry 167, 1479–88 (2010).

26. Duncan, C. E. et al. Prefrontal GABA(A) receptor alpha-subunit expression in normal postnatal human development and schizophrenia. J Psychiatr Res 44, 673–681 (2010).

27. Bosman, L. W. J., Rosahl, T. W. & Brussaard, A. B. Neonatal development of the rat visual cortex: synaptic function of GABAA receptor alpha subunits. J Physiol 545, 169–181 (2002).

28. Katagiri, H., Fagiolini, M. & Hensch, T. K. Optimization of somatic inhibition at critical period onset in mouse visual cortex. Neuron 53, 805–812 (2007).

29. Selemon, L. D. A role for synaptic plasticity in the adolescent development of executive function. Transl Psychiatry 3, e238–e238 (2013).

30. Katz, L. C. & Shatz, C. J. Synaptic activity and the construction of cortical circuits. Science 274, 1133–1138 (1996).

31. Toyoizumi, T. et al. A theory of the transition to critical period plasticity: inhibition selectively suppresses spontaneous activity. Neuron 80, 51–63 (2013).

32. Fagiolini, M. et al. Specific GABAA circuits for visual cortical plasticity. Science 303, 1681–1683 (2004).

33. Ferguson, B. R. & Gao, W.-J. PV Interneurons: Critical Regulators of E/I Balance for Prefrontal Cortex-Dependent Behavior and Psychiatric Disorders. Front Neural Circuits 12, 37 (2018).

34. Cho, R. Y. et al. Development of sensory gamma oscillations and cross-frequency coupling from childhood to early adulthood. Cereb. Cortex 25, 1509–1518 (2015).

35. Buzsáki, G. & Wang, X.-J. Mechanisms of Gamma Oscillations. Annu Rev Neurosci 35, 203–225 (2012).

36. Sohal, V. S. & Rubenstein, J. L. R. Excitation-inhibition balance as a framework for investigating mechanisms in neuropsychiatric disorders. Mol Psychiatry 24, 1248–1257 (2019).

37. Adesnik, H. & Scanziani, M. Lateral competition for cortical space by layer-specific horizontal circuits. Nature 464, 1155–1160 (2010).

38. Yizhar, O. et al. Neocortical excitation/inhibition balance in information processing and social dysfunction. Nature 477, 171–178 (2011).

39. Lally, N. et al. Glutamatergic correlates of gamma-band oscillatory activity during cognition: a concurrent ER-MRS and EEG study. Neuroimage 85 **Pt** **2**, 823–833 (2014).

40. Salinas, E. & Sejnowski, T. J. Correlated neuronal activity and the flow of neural information. Nat.Rev.Neurosci. 2, 539–550 (2001).

41. Mariño, J. et al. Invariant computations in local cortical networks with balanced excitation and inhibition. Nat Neurosci 8, 194–201 (2005).

42. Lim, S. & Goldman, M. S. Balanced cortical microcircuitry for maintaining information in working memory. Nat Neurosci 16, 1306–1314 (2013).

43. Thut, G., Schyns, P. & Gross, J. Entrainment of Perceptually Relevant Brain Oscillations by Non-Invasive Rhythmic Stimulation of the Human Brain. Frontiers in Psychology 2, (2011).

44. Sugiyama, S. et al. The 40-Hz auditory steady-state response enhanced by beta-band subharmonics. Frontiers in Neuroscience 17, (2023).

45. Galambos, R., Makeig, S. & Talmachoff, P. J. A 40-Hz auditory potential recorded from the human scalp. Proceedings of the National Academy of Sciences 78, 2643–2647 (1981).

46. Başar, E., Rosen, B., Başar-Eroglu, C. & Greitschus, F. The associations between 40 Hz-EEG and the middle latency response of the auditory evoked potential. Int J Neurosci 33, 103–117 (1987).

47. Picton, T. W., John, M. S., Dimitrijevic, A. & Purcell, D. Human auditory steady-state responses: Respuestas auditivas de estado estable en humanos. International Journal of Audiology 42, 177–219 (2003).

48. Tada, M. et al. Gamma-Band Auditory Steady-State Response as a Neurophysiological Marker for Excitation and Inhibition Balance: A Review for Understanding Schizophrenia and Other Neuropsychiatric Disorders. Clin EEG Neurosci 51, 234–243 (2020).

49. Sugiyama, S. et al. The Auditory Steady-State Response: Electrophysiological Index for Sensory Processing Dysfunction in Psychiatric Disorders. Front Psychiatry 12, 644541 (2021).

50. Hirano, Y. et al. Spontaneous Gamma Activity in Schizophrenia. JAMA Psychiatry 72, 813–821 (2015).

51. Mathalon, D. H. & Sohal, V. S. Neural Oscillations and Synchrony in Brain Dysfunction and Neuropsychiatric Disorders: It’s About Time. JAMA Psychiatry 72, 840–844 (2015).

52. Uhlhaas, P. J. & Singer, W. Abnormal neural oscillations and synchrony in schizophrenia. Nat.Rev.Neurosci. 11, 100–113 (2010).

53. Thuné, H., Recasens, M. & Uhlhaas, P. J. The 40-Hz Auditory Steady-State Response in Patients With Schizophrenia: A Meta-analysis. JAMA Psychiatry 73, 1145–1153 (2016).

54. O’Donnell, B. F. et al. The auditory steady-state response (ASSR): a translational biomarker for schizophrenia. Suppl Clin Neurophysiol 62, 101–112 (2013).

55. Oda, Y. et al. Gamma band neural synchronization deficits for auditory steady state responses in bipolar disorder patients. PLoS One 7, e39955 (2012).

56. Parker, D. A. et al. Auditory steady-state EEG response across the schizo-bipolar spectrum. Schizophr Res 209, 218–226 (2019).

57. Spencer, K. M., Salisbury, D. F., Shenton, M. E. & McCarley, R. W. Gamma-Band Auditory Steady-State Responses Are Impaired in First Episode Psychosis. Biol Psychiatry 64, 369–375 (2008).

58. Light, G. A. et al. Gamma band oscillations reveal neural network cortical coherence dysfunction in schizophrenia patients. Biol Psychiatry 60, 1231–1240 (2006).

59. Tada, M. et al. Differential Alterations of Auditory Gamma Oscillatory Responses Between Pre-Onset High-Risk Individuals and First-Episode Schizophrenia. Cerebral Cortex 26, 1027–1035 (2016).

60. Parciauskaite, V. et al. 40-Hz auditory steady-state responses and the complex information processing: An exploratory study in healthy young males. PLoS One 14, e0223127 (2019).

61. Tada, M. et al. Alterations of auditory-evoked gamma oscillations are more pronounced than alterations of spontaneous power of gamma oscillation in early stages of schizophrenia. Transl Psychiatry 13, 1–8 (2023).

62. Parciauskaite, V., Bjekic, J. & Griskova-Bulanova, I. Gamma-Range Auditory Steady-State Responses and Cognitive Performance: A Systematic Review. Brain Sci 11, 217 (2021).

63. Vohs, J. L. et al. GABAergic modulation of the 40 Hz auditory steady-state response in a rat model of schizophrenia. International Journal of Neuropsychopharmacology 13, 487–497 (2010).

64. Sivarao, D. V. et al. 40 Hz Auditory Steady-State Response Is a Pharmacodynamic Biomarker for Cortical NMDA Receptors. Neuropsychopharmacol 41, 2232–2240 (2016).

65. Sullivan, E. M., Timi, P., Hong, L. E. & O’Donnell, P. Effects of NMDA and GABA-A Receptor Antagonism on Auditory Steady-State Synchronization in Awake Behaving Rats. International Journal of Neuropsychopharmacology 18, pyu118 (2015).

66. Saunders, J. A., Gandal, M. J. & Siegel, S. J. NMDA antagonists recreate signal-to-noise ratio and timing perturbations present in schizophrenia. Neurobiol Dis 46, 93–100 (2012).

67. Plourde, G., Baribeau, J. & Bonhomme, V. Ketamine increases the amplitude of the 40-Hz auditory steady-state response in humans. Br J Anaesth 78, 524–529 (1997).

68. Farahani, E. D., Wouters, J. & van Wieringen, A. Brain mapping of auditory steady-state responses: A broad view of cortical and subcortical sources. Human Brain Mapping 42, 780–796 (2021).

69. Herdman, A. T. et al. Determination of activation areas in the human auditory cortex by means of synthetic aperture magnetometry. NeuroImage 20, 995–1005 (2003).

70. Popescu, M., Popescu, E.-A., Chan, T., Blunt, S. D. & Lewine, J. D. Spatio-temporal reconstruction of bilateral auditory steady-state responses using MEG beamformers. IEEE Trans Biomed Eng 55, 1092– 1102 (2008).

71. Kuriki, S., Kobayashi, Y., Kobayashi, T., Tanaka, K. & Uchikawa, Y. Steady-state MEG responses elicited by a sequence of amplitude-modulated short tones of different carrier frequencies. Hear Res 296, 25–35 (2013).

72. Farahani, E. D., Goossens, T., Wouters, J. & van Wieringen, A. Spatiotemporal reconstruction of auditory steady-state responses to acoustic amplitude modulations: Potential sources beyond the auditory pathway. NeuroImage 148, 240–253 (2017).

73. Koshiyama, D. et al. Source decomposition of the frontocentral auditory steady-state gamma band response in schizophrenia patients and healthy subjects. Psychiatry and Clinical Neurosciences 75, 172– 179 (2021).

74. Mancini, V. et al. Aberrant Developmental Patterns of Gamma-Band Response and Long-Range Communication Disruption in Youths With 22q11.2 Deletion Syndrome. Am J Psychiatry 179, 204–215 (2022).

75. Tada, M. et al. Global and Parallel Cortical Processing Based on Auditory Gamma Oscillatory Responses in Humans. Cerebral Cortex 31, 4518–4532 (2021).

76. Wang, X. et al. Aberrant Auditory Steady-State Response of Awake Mice After Single Application of the NMDA Receptor Antagonist MK-801 Into the Medial Geniculate Body. International Journal of Neuropsychopharmacology 23, 459–468 (2020).

77. Manting, C. L., Andersen, L. M., Gulyas, B., Ullén, F. & Lundqvist, D. Attentional modulation of the auditory steady-state response across the cortex. NeuroImage 217, 116930 (2020).

78. Shahriari, Y. et al. Impaired auditory evoked potentials and oscillations in frontal and auditory cortex of a schizophrenia mouse model. The World Journal of Biological Psychiatry 17, 439–448 (2016).

79. Ross, B., Hillyard, S. A. & Picton, T. W. Temporal Dynamics of Selective Attention during Dichotic Listening. Cerebral Cortex 20, 1360–1371 (2010).

80. Bouwer, F. L. Neural Entrainment to Auditory Rhythms: Automatic or Top-Down Driven? J. Neurosci. 42, 2146–2148 (2022).

81. Rojas, D. C. et al. Development of the 40Hz steady state auditory evoked magnetic field from ages 5 to 52. Clinical Neurophysiology 117, 110–117 (2006).

82. Poulsen, C., Picton, T. W. & Paus, T. Age-related changes in transient and oscillatory brain responses to auditory stimulation during early adolescence. Dev.Sci. 12, 220–235 (2009).

83. Miskovic, V. et al. Developmental Changes in Spontaneous Electrocortical Activity and Network Organization From Early to Late Childhood. Neuroimage 118, 237–247 (2015).

84. Fagiolini, M. & Hensch, T. K. Inhibitory threshold for critical-period activation in primary visual cortex. Nature 404, 183–186 (2000).

85. McKeon, S. D. et al. Aperiodic EEG and 7T MRSI evidence for maturation of E/I balance supporting the development of working memory through adolescence. Developmental Cognitive Neuroscience 66, 101373 (2024).

86. Perica, M. I. et al. Development of frontal GABA and glutamate supports excitation/inhibition balance from adolescence into adulthood. Progress in Neurobiology 219, 102370 (2022).

87. Delorme, A. & Makeig, S. EEGLAB: an open source toolbox for analysis of single-trial EEG dynamics including independent component analysis. J. Neurosci. Methods 134, 9–21 (2004).

88. Pan, J. w., Avdievich, N. & Hetherington, H. p. J-refocused coherence transfer spectroscopic imaging at 7 T in human brain. Magnetic Resonance in Medicine 64, 1237–1246 (2010).

89. Hetherington, H. P., Avdievich, N. I., Kuznetsov, A. M. & Pan, J. W. RF shimming for spectroscopic localization in the human brain at 7 T. Magnetic Resonance in Medicine 63, 9–19 (2010).

90. David, O., Kilner, J. M. & Friston, K. J. Mechanisms of evoked and induced responses in MEG/EEG. NeuroImage 31, 1580–1591 (2006).

91. Hirano, Y. et al. Auditory Cortex Volume and Gamma Oscillation Abnormalities in Schizophrenia. Clin EEG Neurosci 51, 244–251 (2020).

92. Grent-’t-Jong, T., Brickwedde, M., Metzner, C. & Uhlhaas, P. J. 40-Hz Auditory Steady-State Responses in Schizophrenia: Toward a Mechanistic Biomarker for Circuit Dysfunctions and Early Detection and Diagnosis. Biological Psychiatry 94, 550–560 (2023).

93. Spencer, K. M., Niznikiewicz, M. A., Nestor, P. G., Shenton, M. E. & McCarley, R. W. Left auditory cortex gamma synchronization and auditory hallucination symptoms in schizophrenia. BMC Neurosci 10, 85 (2009).

94. Provencher, S. W. Automatic quantitation of localized in vivo 1H spectra with LCModel. NMR Biomed 14, 260–264 (2001).

95. Rideaux, R. No balance between glutamate+glutamine and GABA+ in visual or motor cortices of the human brain: A magnetic resonance spectroscopy study. NeuroImage 237, 118191 (2021).

96. Steel, A., Mikkelsen, M., Edden, R. A. E. & Robertson, C. E. Regional balance between glutamate+glutamine and GABA+ in the resting human brain. Neuroimage 220, 117112 (2020).

97. Wood, S. N. *Generalized Additive Models: An Introduction with R*. (Chapman and Hall/CRC, New York, 2017). doi:10.1201/9781315370279.

98. Luna, B., Velanova, K. & Geier, C. F. Development of eye-movement control. Brain Cogn 68, 293–308 (2008).

99. McKeon, S. D. et al. Aperiodic EEG and 7T MRSI evidence for maturation of E/I balance supporting the development of working memory through adolescence. 2023.09.06.556453 Preprint at 10.1101/2023.09.06.556453 (2023).

100. Buuren, S. van & Groothuis-Oudshoorn, K. mice: Multivariate Imputation by Chained Equations in R. Journal of Statistical Software 45, 1–67 (2011).

101. Wood, S. N. A simple test for random effects in regression models. Biometrika 100, 1005–1010 (2013).

102. Bates, D., Mächler, M., Bolker, B. & Walker, S. Fitting Linear Mixed-Effects Models Using lme4. Journal of Statistical Software 67, 1–48 (2015).

103. Larsen, B. et al. A developmental reduction of the excitation:inhibition ratio in association cortex during adolescence. Science Advances 8, eabj8750 (2022).

104. Lewis, D. A., Hashimoto, T. & Volk, D. W. Cortical inhibitory neurons and schizophrenia. Nat Rev Neurosci 6, 312–324 (2005).

105. Gonzalez-Burgos, G. & Lewis, D. A. GABA neurons and the mechanisms of network oscillations: implications for understanding cortical dysfunction in schizophrenia. Schizophr.Bull. 34, 944–961 (2008).

106. Erickson, S. L. & Lewis, D. A. Postnatal development of parvalbumin- and GABA transporter-immunoreactive axon terminals in monkey prefrontal cortex. Journal of Comparative Neurology 448, 186– 202 (2002).

107. Hoftman, G. D. & Lewis, D. A. Postnatal Developmental Trajectories of Neural Circuits in the Primate Prefrontal Cortex: Identifying Sensitive Periods for Vulnerability to Schizophrenia. Schizophr Bull 37, 493– 503 (2011).

108. Cho, R. Y. et al. Development of Sensory Gamma Oscillations and Cross-Frequency Coupling from Childhood to Early Adulthood. Cereb Cortex 25, 1509–1518 (2015).

109. Huttenlocher, P. R. & Dabholkar, A. S. Regional differences in synaptogenesis in human cerebral cortex. J Comp Neurol 387, 167–178 (1997).

110. Wainio-Theberge, S., Wolff, A. & Northoff, G. Dynamic relationships between spontaneous and evoked electrophysiological activity. Commun Biol 4, 1–17 (2021).

111. Myers, N. E., Stokes, M. G., Walther, L. & Nobre, A. C. Oscillatory Brain State Predicts Variability in Working Memory. J Neurosci 34, 7735–7743 (2014).

112. Whitford, T. J. et al. Brain maturation in adolescence: Concurrent changes in neuroanatomy and neurophysiology. Human Brain Mapping 28, 228–237 (2007).

113. Tierney, A., Strait, D. L., O’Connell, S. & Kraus, N. Developmental changes in resting gamma power from age three to adulthood. Clinical Neurophysiology 124, 1040–1042 (2013).

114. Anderson, A. J. & Perone, S. Developmental change in the resting state electroencephalogram: Insights into cognition and the brain. Brain Cogn 126, 40–52 (2018).

115. Matousek, M. & Petersén, I. Automatic evaluation of EEG background activity by means of age-dependent EEG quotients. Electroencephalogr Clin Neurophysiol 35, 603–612 (1973).

116. Carlén, M. et al. A critical role for NMDA receptors in parvalbumin interneurons for gamma rhythm induction and behavior. Mol Psychiatry 17, 537–548 (2012).

117. Buzsáki, G., Anastassiou, C. A. & Koch, C. The origin of extracellular fields and currents--EEG, ECoG, LFP and spikes. Nat Rev Neurosci 13, 407–420 (2012).

118. Sohal, V. S., Zhang, F., Yizhar, O. & Deisseroth, K. Parvalbumin neurons and gamma rhythms enhance cortical circuit performance. Nature 459, 698–702 (2009).

119. Burt, J. B. et al. Hierarchy of transcriptomic specialization across human cortex captured by structural neuroimaging topography. Nature Neuroscience 21, 1251 (2018).

120. Wang, H., Stradtman, G. G., Wang, X.-J. & Gao, W.-J. A specialized NMDA receptor function in layer 5 recurrent microcircuitry of the adult rat prefrontal cortex. Proceedings of the National Academy of Sciences 105, 16791–16796 (2008).

121. Erisir, A. & Harris, J. L. Decline of the critical period of visual plasticity is concurrent with the reduction of NR2B subunit of the synaptic NMDA receptor in layer 4. J. Neurosci. 23, 5208–5218 (2003).

122. Sydnor, V. J. et al. Neurodevelopment of the association cortices: Patterns, mechanisms, and implications for psychopathology. Neuron 109, 2820–2846 (2021).

123. Hilgetag, C. C., Beul, S. F., van Albada, S. J. & Goulas, A. An architectonic type principle integrates macroscopic cortico-cortical connections with intrinsic cortical circuits of the primate brain. Netw Neurosci 3, 905–923 (2019).

124. Simmonds, D. J., Hallquist, M. N., Asato, M. & Luna, B. Developmental stages and sex differences of white matter and behavioral development through adolescence: a longitudinal diffusion tensor imaging (DTI) study. NeuroImage 92, 356–368 (2014).

125. Tervo-Clemmens, B., et al. A Canonical Trajectory of Executive Function Maturation During the Transition from Adolescence to Adulthood. Preprint at 10.31234/osf.io/73yfv (2022).

126. Ravindranath, O., Calabro, F. J., Foran, W. & Luna, B. Pubertal development underlies optimization of inhibitory control through specialization of ventrolateral prefrontal cortex. Developmental Cognitive Neuroscience 58, 101162 (2022).

127. Goldman-Rakic, P. S. Architecture of the Prefrontal Cortex and the Central Executive. Annals of the New York Academy of Sciences 769, 71–84 (1995).

128. Buschman, T. J. Balancing Flexibility and Interference in Working Memory. Annu Rev Vis Sci 7, 367– 388 (2021).

129. Luna, B., Marek, S., Larsen, B., Tervo-Clemmens, B. & Chahal, R. An Integrative Model of the Maturation of Cognitive Control. Annu Rev Neurosci 38, 151–170 (2015).

130. Blakemore, S.-J. & Mills, K. L. Is adolescence a sensitive period for sociocultural processing? Annu Rev Psychol 65, 187–207 (2014).

131. Steinberg, L. A social neuroscience perspective on adolescent risk-taking. Dev Rev 28, 78–106 (2008).

132. Chavez-Baldini, U. et al. The relationship between cognitive functioning and psychopathology in patients with psychiatric disorders: a transdiagnostic network analysis. Psychol Med 53, 476–485.

133. van Bueren, N. E. R., van der Ven, S. H. G., Hochman, S., Sella, F. & Cohen Kadosh, R. Human neuronal excitation/inhibition balance explains and predicts neurostimulation induced learning benefits. PLoS Biol 21, e3002193 (2023).

134. Selten, M., van Bokhoven, H. & Nadif Kasri, N. Inhibitory control of the excitatory/inhibitory balance in psychiatric disorders. F1000Res 7, (2018).

135. Gonzalez-Burgos, G., Hashimoto, T. & Lewis, D. A. Alterations of cortical GABA neurons and network oscillations in schizophrenia. Curr Psychiatry Rep 12, 335–344 (2010).

