## Supplement for "Prefrontal Excitation/Inhibition Balance Supports Adolescent Enhancements in Circuit Signal to Noise Ratio"

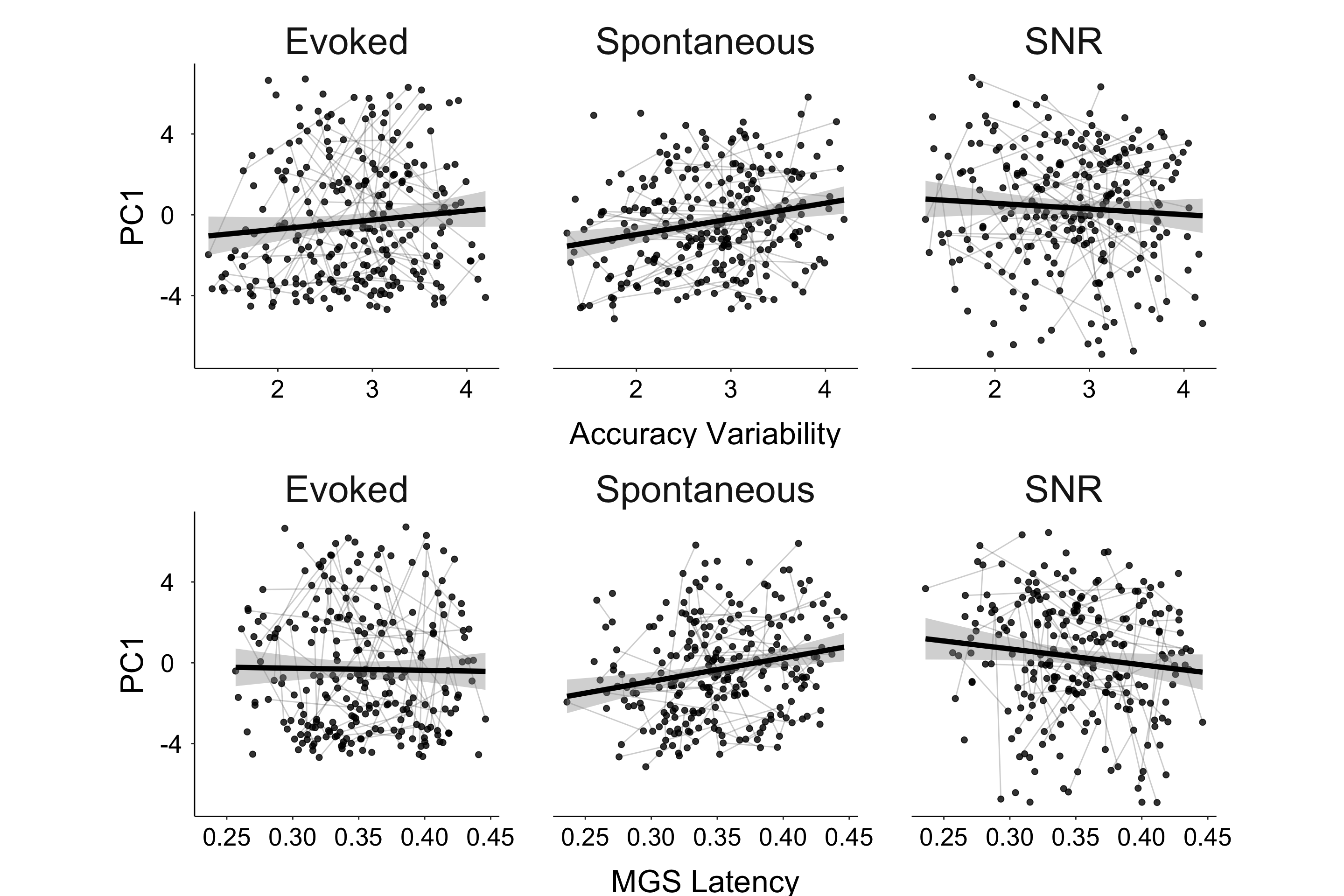

Supplement Figure 1. Auditory Measures vs Behavioral Measures.

Top: Evoked (left), spontaneous (middle), and cortical SNR (right) vs accuracy trial variability. Bottom: Evoked (left), spontaneous (middle), and cortical SNR (right) vs response latency.

Supplement Table 1. Auditory Measures vs Behavioral Measures

|  |  | **Evoked** | | | **Spontaneous** | | | **Cortical SNR** | | |
| --- | --- | --- | --- | --- | --- | --- | --- | --- | --- | --- |
|  |  | **b** | **t** | **p** | **b** | **t** | **p** | **b** | **t** | **p** |
| **Behavioral main effects** | **Accuracy** | -0.24 | -1.09 | 0.27 | 0.21 | 1.4 | 0.16 | **-0.53** | **-2.56** | **0.01** |
|  | **Accuracy Var** | 0.22 | 0.74 | 0.45 | 0.30 | 1.49 | 0.13 | -0.28 | -1.01 | 0.31 |
|  | **Latency** | -4.23 | -0.88 | 0.37 | 5.98 | 1.88 | 0.06 | -6.65 | -1.46 | 0.14 |
|  | **Latency. Var** | -9.29 | -1.19 | 0.23 | **14.76** | **2.85** | **0.004** | **-16.3** | **-2.2** | **0.02** |
|  |  | **b** | **t** | **p** | **b** | **t** | **p** | **b** | **t** | **p** |
| **Behavioral age interaction** | **Accuracy** | -0.02 | 0.50 | 0.61 | 0.01 | 0.27 | 0.78 | -0.01 | -0.33 | 0.74 |
|  | **Accuracy Var** | 0.01 | 0.21 | 0.83 | 0.02 | 0.57 | 0.56 | -0.02 | -0.49 | 0.62 |
|  | **Latency** | 0.35 | 0.41 | 0.68 | -0.13 | -0.23 | 0.81 | 0.33 | 0.40 | 0.68 |
|  | **Latency. Var** | 0.53 | 0.422 | 0.67 | 0.93 | 1.07 | 0.28 | -0.47 | -0.38 | 0.69 |
